# Infant neural sensitivity to affective touch is associated with maternal postpartum depression

**DOI:** 10.1101/2024.02.21.581204

**Authors:** Madelyn G. Nance, Zackary T. Landsman, Gregory J Gerling, Meghan H. Puglia

## Abstract

Classic attachment theory emphasizes the sensitivity of the parent to perceive and appropriately respond to the infant’s cues. However, parent-child attachment is a dyadic interaction that is also dependent upon the sensitivity of the child to the early caregiving environment. Individual differences in infant sensitivity to parental cues is likely shaped by both the early caregiving environment as well as the infant’s neurobiology, such as perceptual sensitivity to social stimuli. Here, we investigated associations between maternal postpartum depression and infant neurological sensitivity to affective touch using brain signal entropy – a metric of the brain’s moment-to-moment variability related to signal processing. We recruited two independent samples of infants aged 0-5 months. In Sample 1 (n=79), we found increased levels of maternal postpartum depression were associated with diminished perceptual sensitivity – i.e. lower entropy – to affective tactile stimulation specifically within the primary somatosensory cortex. In Sample 2 (n=36), we replicated this finding and showed that this effect was not related to characteristics of the touch administered during the experiment. These results suggest that decreased affective touch early in life – a common consequence of postpartum depression – likely impacts the infant’s perceptual sensitivity to affective touch and ultimately the formation of experience-dependent neural networks that support the successful formation of attachment relationships.

## 2. Introduction

Somatosensation is the first sensory system to mature (Gao, Alcauter, Elton, et al., 2015; Gao, Alcauter, Smith, et al., 2015), allowing the baby to integrate signals from their external environment with internal processes. Touch is therefore one of the earliest ways that mothers communicate with their infants, often by providing loving, affective touch. Affective touch is a form of social touch that preferentially activates C-tactile afferents – a mechanoreceptor found on hairy skin that maximally responds to slow, gentle stroking (Olausson et al., 2010; Vallbo et al., 1999) from social partners (Aguirre et al., 2019; Mantis et al., 2019; Pawling et al., 2017; Sailer & Leknes, 2022). The primary somatosensory cortex (S1) is a region of the brain responsible for processing tactile information including discriminating the affective content of touch (Suvilehto et al., 2021). While many other socio-emotional brain regions are also associated with affective touch processing in older children and adults (Bjornsdotter et al., 2014; Croy et al., 2019; Gordon et al., 2013; McGlone et al., 2017), there is significant evidence that infants have not yet developed this network and primarily discriminate between affective and non-affective touch in S1 (Miguel, Gonçalves, et al., 2019; Miguel, Lisboa, et al., 2019; Pirazzoli et al., 2019; c.f. Jönsson et al., 2018).

Affective touch serves as an early avenue for establishing mother-infant bonds. The bond between mother and infant shapes social and neural development and has important implications for later attachment styles (Benoit, 2004; Bowlby, 1982; Deave et al., 2008; Toth et al., 2009). Strong, secure attachment to the mother enables the child to engage with the environment and form social bonds with others (Ainsworth & Bell, 1970; Rajecki & Obmascher, 1978). Attachment theory asserts that physical contact is the most important signal to the infant that they are in a secure relationship (Duhn, 2010; Reite, 1990; Weiss et al., 2000). It has repeatedly been demonstrated that increased physical contact in infancy, specifically increased affective touch (Weiss et al., 2000) is associated with increased secure attachment between mother and infant (Ainswoth, 1967; Anisfeld et al., 1990). For example, in a particularly striking illustration of the importance of touch in development, Harlow demonstrated that infant monkeys preferentially spend time with a surrogate mother that provides comforting tactile stimulation over one which provides nutrition (Harlow & Zimmermann, 1959).

The mother-infant attachment bond is highly sensitive to the mother’s psychological wellbeing which enables her to successfully perceive, interpret, and respond to her infant’s cues (Edhborg et al., 2011; Moehler et al., 2006; O’Higgins et al., 2013). Postpartum depression (PPD) is the most common complication of birth in the United States and causes a sad or depressed mood, feelings of guilt, and/or a lack of interest in the new baby that impairs mother-infant bonding (Deligiannidis et al., 2023; Toohey, 2012). Decreased communication and social interaction between mother and baby are common with PPD, alter the early life environment, and put infants of depressed mothers at an increased risk of delays in cognitive and motor development (Field, 2010). PPD also has significant effects on the frequency and forms of touch that mothers use towards their infants (Ferber et al., 2008). For example, higher scores on PPD screening measures are correlated with a lower frequency and duration of playful touch (Mantis et al., 2019) and increased use of negative or assertive forms of touch such as poking or pulling and intrusively interrupting their baby from playing (Herrera et al., 2004; Malphurs et al., 1996). PPD is thus a considerable obstacle to forming a dyadic, responsive, and secure infant-mother attachment.

In a contemporaneous study, Shekhar et al. (Shekhar et al., 2024) investigated associations between prenatal depression symptoms at 34 weeks’ gestation and neural response during affective touch in a small sample of two-year-olds using functional near infrared spectroscopy (fNIRS). This study found that increased prenatal depression symptoms were associated with decreased activation of S1 during affective touch, specifically. These results contrast another recent fNIRS study that assessed maternal sensitivity at 7 months of age and neural response to affective touch at 12 months of age (Mateus et al., 2021). This study found decreased maternal sensitivity was associated with increased neural response to affective touch within S1. While these studies indicate that maternal behavior such as depression and sensitivity modulate infants’ neural response to affective touch in S1, the primary limitations to this prior work are the large gap between assessments of maternal behavior and infant neural response to affective touch, measurement in older infants/toddlers, small sample sizes, and the use of NIRS, which is an indirect measure of brain activity based on the blood oxygen level dependent response.

Here, we investigated the impacts of maternal depression on neural response to affective touch using electroencephalography (EEG) in two independent samples in early infancy – a critical period for attachment formation. Classic attachment theory emphasizes the importance of the mother’s ability to provide affective touch in response to the infant’s socioemotional cues (Chris Fraley, 2018; Rajecki & Obmascher, 1978), but mother-infant attachment arises from a dyadic interaction that is also dependent upon the infant’s perceptual sensitivity to the mother’s nurturing touch. It is increasingly understood that perceptual sensitivity is related to signal variability in the brain (Garrett et al., 2013; Puglia et al., 2020; Ward et al., 2006). This neural variability facilitates signal detection, synchrony between neurons, and the formation of experience-dependent neural networks associated with the development of perceptual and cognitive stability (Puglia et al., 2020; Garrett et al., 2011; Malins et al., 2018; McIntosh et al., 2008, 2010; Waschke et al., 2017). One way to quantify such brain signal variability is through the use of multiscale entropy (Costa et al., 2002; Courtiol et al., 2016). Entropy is a measure of the randomness or unpredictability of a signal. Complex biological signals – such as the electrical signaling in the brain – consist of both random and deterministic properties across multiple time scales that can be measured via EEG (Ward et al., 2006). To quantify multiscalar variability, entropy can be computed across multiple scales by coarse graining the time series (Costa et al., 2002). Here we use multiscale (scale-wise) entropy (Puglia et al., 2022) to assess how perceptual sensitivity to affective touch varies as a function of the early caregiving environment. We hypothesized that infants whose mothers had more PPD symptoms would display decreased levels of brain signal entropy during affective touch in S1, suggesting decreased perceptual sensitivity for this form of touch.

## 3. Methods

### 3.1 Participants

Two samples of infant-mother dyads participated in the present study to determine the effect of maternal PPD on infant brain signal entropy during affective touch. The primary caregiver accompanied the infant to the study appointment and provided written informed consent for a protocol approved by the University of Virginia (UVA) Health Sciences Research Institutional Review Board (HSR19514, PI: Puglia). All procedures were performed in compliance with relevant laws and institutional guidelines. Families were paid $50 for their participation. Only families in which the birth mother lived with the child were included in the study.

The first sample (Sample 1) consisted of 79 infants (45 F) and their mothers recruited through the UVA Hospital (n=67) and the greater Charlottesville area (n=12). Infants ranged in age from 12 to 169 (M=65.65, SD=42.99) days. Mothers ranged in age from 21 to 45 (M=31.53, SD=4.45) years. A second independent sample (Sample 2) of 36 infants (14 F) and their mothers were recruited through the UVA Hospital to replicate and extend our findings. Infants ranged in age from 7 to 147 (M=76.06, SD=44.62) days. Mothers ranged in age from 22 to 40 (M=32.56, SD=5.26) years. A post hoc power analysis with the simr package (Green & MacLeod, 2016) in R v4.3.1 (R Core Team, 2020) revealed that we achieved 97% power [95% confidence interval (CI): 91.48-99.38] in Sample 1 to detect the effect of maternal PPD on infant brain signal entropy, and would require 40 participants to achieve 80% power to detect this effect in the replication sample. After data preprocessing (see below), there were 36 participants in Sample 2 with usable EEG data, and we achieved 73% power [95% CI: 63.20-81.39].

There was no significant difference in infant age (t(113) = -1.19, p=.237) or mother age ((t(113) = -1.08, p=.283) across the two samples. Both samples have a racial and ethnic makeup reflective of the greater Charlottesville area (*U.S. Census Bureau QuickFacts*, n.d.). Demographic characteristics are detailed in Table 1.

**Table 1:** Participant Demographics. Demographic information was extracted from the electronic health record or self-reported at the study visit.

| Demographic |  | Sample 1 (n=79) | Sample 2 (n=36) |
| --- | --- | --- | --- |
| Mother's Age (M[SD]) |  | 31.53 [4.45] | 32.56 [5.26] |
| Total Household Income (M[SD]) | | \$110,075.50<br>[\$66,766.43] | \$92,459.29<br>[\$58,641.19] |
| Delivery Method (n) | Vaginal | 56 | 23 |
|  | C-section | 23 | 13 |
| Mother's Race (n) | Asian | 4 | 2 |
|  | Black or African American | 11 | 4 |
|  | White | 60 | 29 |
|  | Other | 4 | 1 |
| Mother's Ethnicity (n) | Not Spanish / Hispanic | 75 | 30 |
|  | Spanish/ Hispanic | 4 | 6 |
| Mother's Marital Status (n) | Single | 14 | 6 |
|  | Married | 62 | 29 |
|  | Unknown | 3 | 1 |
| Mother's Education Level (n) | Grade School | 1 | 1 |
|  | High School Graduate | 13 | 6 |
|  | Associate's Degree | 4 | 2 |
|  | Bachelor's Degree | 26 | 13 |
|  | Master's Degree | 17 | 10 |
|  | Doctorate / Professional Degree | 15 | 3 |
|  | Unknown | 3 | 1 |

### 3.2 Edinburgh Postnatal Depression Scale

The Edinburgh Postnatal Depression Scale (EPDS) is the most common screening tool used to identify PPD (Lubotzky-Gete et al., 2021) and produces scores that are stable throughout the extent of the postpartum period (Abdollahi et al., 2017; Kubota et al., 2018). This scale offers enhanced specificity for the detection of PPD relative to traditional depression questionnaires that focus heavily on somatic symptoms of depression such as lethargy, which is often unrelated to depression in the postpartum period (Cox et al., 1987).

For the present study, EPDS scores were extracted from the birth mother’s medical chart using the UVA hospital’s Electronic Health Records (EHR) system or, if unavailable via EHR, through completion of a questionnaire during the study appointment. When the EHR contained multiple occurrences of EPDS, the score closest to the child’s EEG date was selected. On average, the lag between the completion of EPDS and EEG collection was 39.15 days for Sample 1 and 41.72 days for Sample 2. There was no significant difference in days between EPDS and EEG collection across samples (t(113) = - 0.35, p=.731).

EPDS scores ranged from 0 to 17 (*M*=4.73, *SD*=4.31) in Sample 1, and from 0 to 16 (*M*=5.92, *SD*=4.49) in Sample 2. EPDS scores did not significantly vary across samples (*t*(113) = -1.35, p=.181). Twenty mothers in Sample 1, and 15 mothers in Sample 2 exceeded the EPDS cutoff score of 7 (McCabe-Beane et al., 2016) for postpartum depression. Associations between demographic characteristics and EPDS scores are detailed in the Supplemental Materials.

### 3.3 Affective Touch Paradigm

Infants participated in a tactile stimulation EEG paradigm designed to activate C-tactile afferents. In this paradigm, an experimenter blind to the mother’s EPDS score administered brushstrokes to the infant’s forearm in the proximal to distal direction with a soft bristled brush at 3 cm/s – a rate that maximally activates C-tactile afferents and reflects the way a mother typically caresses her infant (Pawling et al., 2017). In the affective touch condition, infants were stroked directly on the skin of their forearm (Figure 1A). In the non-affective touch sensorimotor control condition, the experimenter placed a thin medical-grade plastic film over the infants’ skin and delivered the same 3 cm/s stroke to the infant’s forearm through the plastic film (Figure 1B). As C-tactile afferents are activated via mechanoreceptors on hairy skin, the plastic film effectively blocks C-tactile activation (Rezaei et al., 2021) while allowing all other touch characteristics including stroking rate and location to remain identical across conditions. Prior to beginning the paradigm, a 3 cm mark was made along the infant’s forearm with a skin-safe marker. The experimenter was aided in accurately timing the touch through a 1-s metronome presented to the experimenter via headphones. The order of conditions was randomized across participants and repeated in a block design consisting of eighteen 1-s strokes per block. The paradigm continued until a maximum of 8 blocks of each condition were completed or until the infant was no longer compliant.

**Figure 1:**
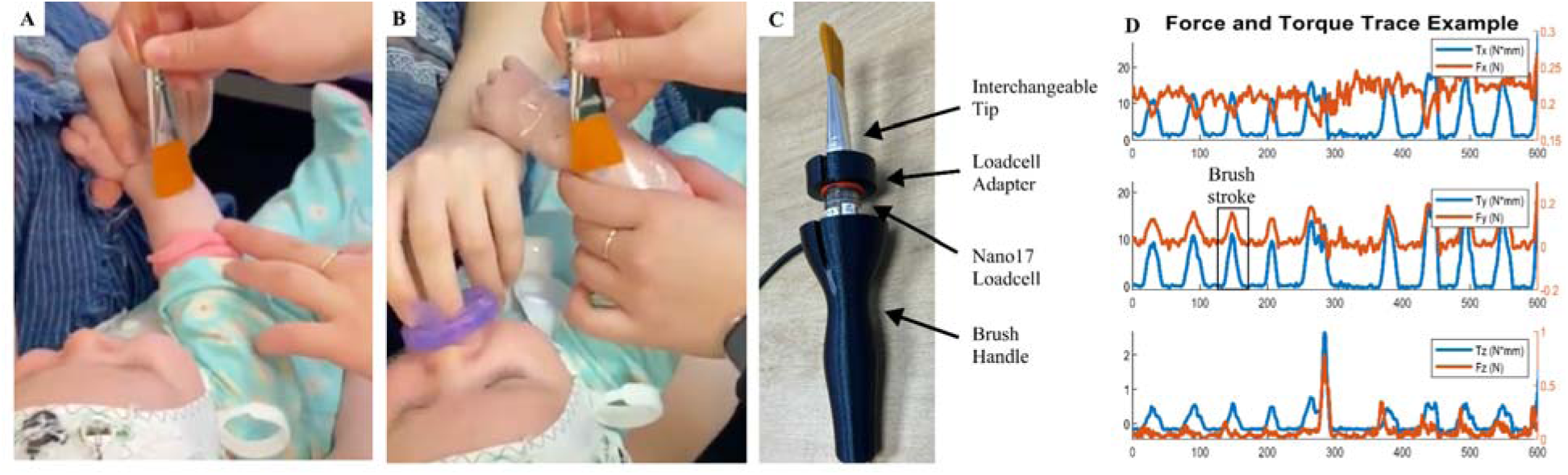
Affective touch paradigm and application of engineered brush. **(A)** Brushing was administered with a soft-tipped brush onto direct skin to elicit C-tactile afferent activation (affective touch) and (**B)** over a layer of plastic film (non-affective touch). **(C)** For Sample 2 subjects, the brushing delivered by each research assistant was recorded using an engineered brush to capture 6-axis force and torque. **(D)** The measurement of torque (blue) was used to quantify the brushing applied to each participant due to the measure’s robustness to noise relative to force (orange).

To establish whether brushing techniques differed as a function of condition, EPDS, or entropy, Sample 2 underwent the same affective touch paradigm with an engineered brush developed by our team to capture 6-axis force and torque of each brushstroke using a Nano-17 load cell (Nano17, ATI Industrial Automation, Apex, NC) housed in a 3D printed frame (Figure 1C & D) (Landsman et al., 2023).

### 3.4 Brushstroke Feature Extraction

Brushing application was quantified by torque magnitude through the aggregation of the torque x, y, and z components. Torque is the rotational equivalent of linear force; in this work it can be thought of as the bending of the brush bristles. Similar to force, more pressure applied during brushing results in more torque generated. The measurement of torque magnitude was used in this work due to its robustness to noise in dynamic human interactions, whereas force is very sensitive to inertia and gravity due to orientation (Figure 1D) (Landsman et al., 2023). The mean torque magnitude of each brushstroke was analyzed using a contact detection algorithm. Across 397 recorded experiment blocks, 8 blocks were removed due to collection error. The feature extraction of each block consisted of identifying individual brushstrokes after detecting contact periods. The contact detection algorithm first tared each component of torque using an end period of “no contact” in each brushing file. Then contact was determined by locating local maxima and minima in the torque magnitude aggregate, thus finding the onset and offloading of torque for each brushstroke. Each brushstroke endpoint was then adjusted to a contact threshold of 1.5 N*mm. The analysis detected 6,151 brushstrokes, from which mean torque magnitude was extracted during each individual brushstroke. Error brush strokes were identified in the set as those with a duration greater than 2 seconds or containing a negative magnitude mean, resulting in the removal of 23 brushstrokes. Brushstroke data for an additional baby was not captured due to recording error. Outlier brush strokes were then identified for each subject as those that had a torque magnitude more than 3 median absolute deviations above or below the median torque magnitude value, resulting in the removal of 98 brushstrokes.

### 3.5 EEG acquisition and analysis

EEG data was collected from 105 participants for Sample 1 and 44 participants for Sample 2. EEG was recorded using 32 Ag/AgCl active actiCap slim electrodes (Brain Products GmbH, Germany) affixed to an elastic cap using the 10–20 electrode placement (Figure 2A). EEG was amplified with a BrainAmp DC Amplifier and digitized using BrainVision Recorder software with a sampling rate of 5000 Hz, online referenced to FCz, and online band-pass filtered between 0.1 and 1000 Hz. Data preprocessing and entropy calculation were accomplished using the Automated Preprocessing Pipe-Line for the Estimation of Scale-wise Entropy from EEG Data (Puglia et al., 2022) implemented with EEGLab v2022.1 (Delorme & Makeig, 2004) in MATLAB v2020a (The MathWorks Inc., 2022). Data were first downsampled to 250 Hz, band-pass filtered from 0.3-50 Hz, and segmented into 1-s epochs time locked to the onset of each brush stroke. On average, infants in Sample 1 received 117.11 affective and 116.62 nNon-affective strokes. Infants in Sample 2 received 108.82 affective and 109.23 non-affective strokes on average. Epochs contaminated with extreme artifacts (>= 500 µV) were removed, and the data then underwent independent component analysis. Problematic components were identified and removed using the adjusted_ADJUST algorithm (Leach et al., 2020). On average, 4.92 components for Sample 1 and 5.12 components for Sample 2 were removed. Then, epochs in which the standard deviation exceeded 80 µV within a 200-ms sliding window with a 100-ms window step were discarded as artifacts, and channels identified as problematic via the FASTER algorithm (Nolan et al., 2010) were interpolated. The number of trials removed from the affective touch (M=48.47) and non-affective touch (M=46.40) conditions did not differ (t(314)=0.53, p=.595). On average 0.86 channels for Sample 1 and 0.98 channels for Sample 2 were interpolated. Participants with fewer than 10 artifact-free trials per condition after preprocessing were removed (Sample 1: n=26 excluded, n=79 retained; Sample 2: n=8 excluded, n=36 retained). Cleaned EEG data were then referenced to the average of all scalp electrodes. We selected the 10 affective touch and 10 non-affective touch epochs with a total global field power (GFP) (McIntosh et al., 2008) closest to the median GFP for each participant for inclusion in the entropy calculation. Finally, multiscale (scale-wise) entropy was computed on these trials using APPLESEED with a similarity criterion r=0.5, and a pattern length m=2. Entropy values were averaged across scales and channels to create two brain regions of interest – primary somatosensory cortex (S1) recorded from C3 and C4 electrodes (Figure 2A), and the whole brain excluding S1 (WB) recorded from all other scalp electrodes. Average entropy curves for each condition and brain region are plotted in Figure 2B.

**Figure 2.**
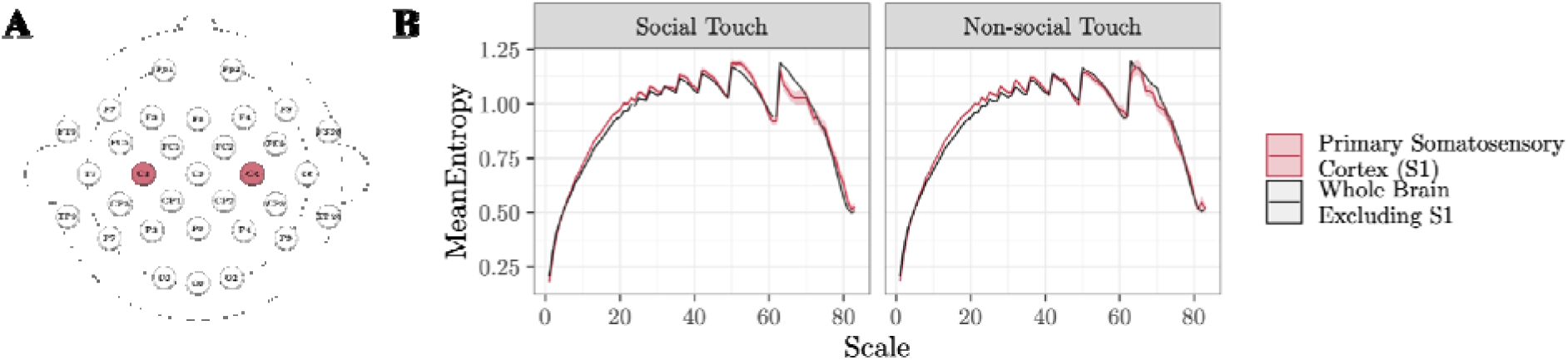
EEG montage and entropy curves. **(A)** Electroencephalography (EEG) was recorded from 32 scalp electrodes. Brain regions of interest included the primary somatosensory cortex (S1), an average of electrodes C3 and C4 (red), and the whole brain excluding S1, the average of all other scalp electrodes (white). **(B)** Multiscale (scale-wise) entropy was computed for each brain region and experimental condition.

### 3.6 Statistical Analysis

We employed a linear mixed effects model in R using the lmerTest package (Kuznetsova et al., 2017) to test for associations between EPDS and brain signal entropy across experimental conditions (affective touch; non-affective touch) and brain regions of interest (S1; WB). All variables were first standardized, then interactions between condition, brain region, and EPDS scores were entered into the model as independent variables to predict brain signal entropy.

In subsequent exploratory analyses, we investigated whether these same associations varied across EEG frequency bands. Separate linear mixed effects models were run as above with brain signal entropy averaged across scales in the following frequency bands: Delta (0-3 Hz; scales 63-83), Theta (4-7 Hz; scales 32-62), Alpha (8-12 Hz; scales 20-31), Beta (13-29 Hz; scales 9-19), and Gamma (30+ Hz; scales 1-8).

Sample 2 underwent the affective touch paradigm using an engineered brush. To test for differences in how these infants were touched as a function of task condition, we computed a two-sample t-test on mean torque magnitude. Additionally, we assessed whether there were differences in how babies were touched during the affective touch condition as a function of mother’s EPDS score via a linear model predicting EPDS scores from standardized mean torque magnitude. Finally, we employed four linear models to test for associations between standardized mean torque magnitude and brain signal entropy for each condition (affective touch; non-affective touch) and brain region (WB; S1).

### 3.7 Data availability statement

Data will be made available upon reasonable request.

## 4. Results

We ran a linear mixed effects model including touch condition (affective; non-affective), brain region of interest (S1; WB), EPDS score, and the interactions among these terms as independent variables and brain signal entropy as the dependent variable. In Sample 1, we found the slope between EPDS and brain signal entropy during affective touch within S1 significantly differed from the slope between EPDS and brain signal entropy during affective touch within WB (β = 0.30, p = .028), non-affective touch within S1 (β = 0.42, p = .002), and non-affective touch within WB (β = 0.42, p = .002, Figure 3). In Sample 2, we replicated this finding showing that the slope between EPDS and brain signal entropy during affective touch within S1 significantly differed from the slope between EPDS and brain signal entropy during affective touch within WB (β **=** 0.36, p = .020), non-affective touch within S1 (β = 0.47, p = .005), and non-affective touch within WB (β = 0.38, p = .020). An exploratory post-hoc analysis revealed that these associations between EPDS and brain signal entropy were consistent regardless of infant age, but strongest in younger infants (Supplemental Materials).

**Figure 3.**
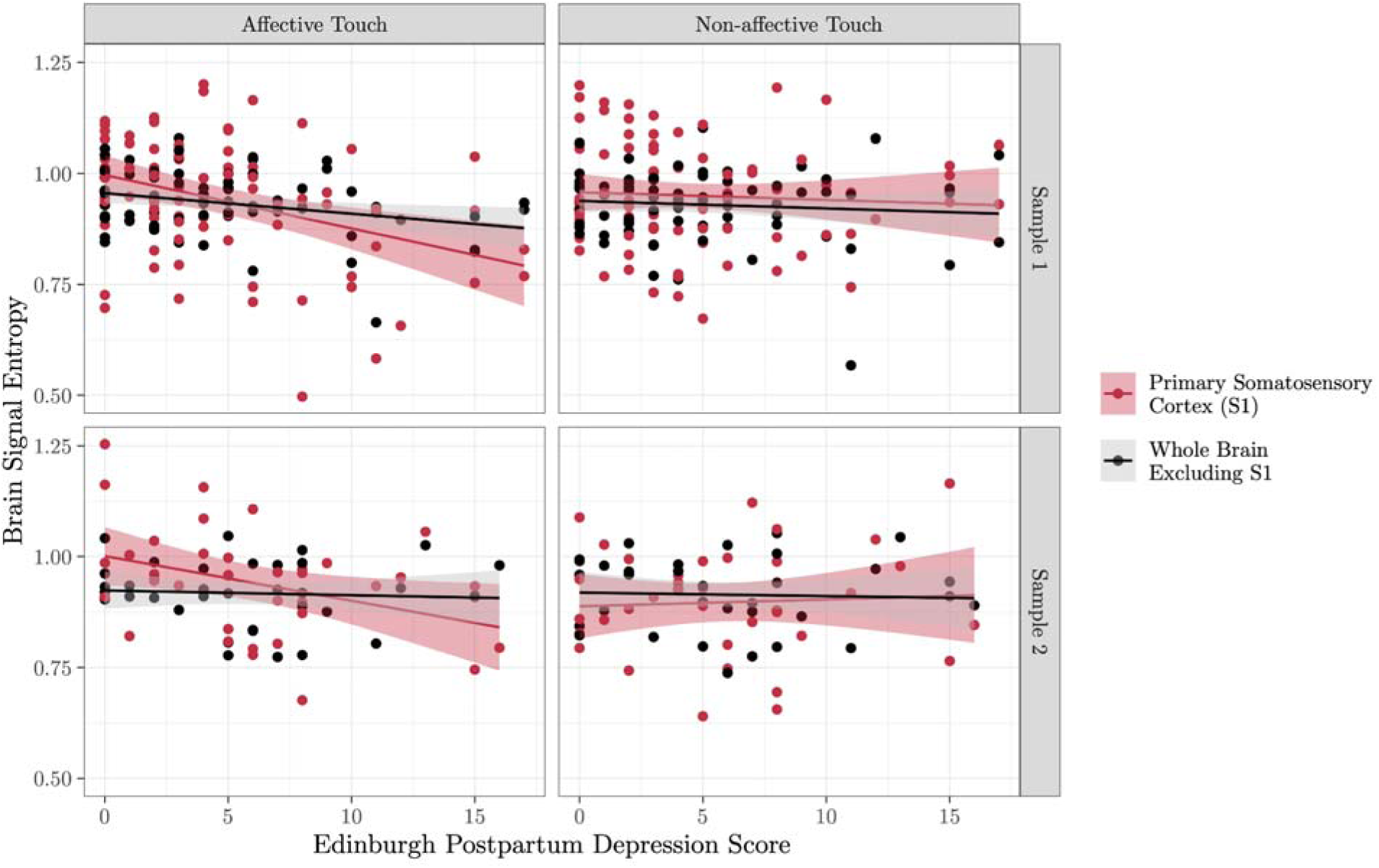
Brain signal entropy during affective touch is associated with maternal postpartum depression in primary somatosensory cortex. A linear mixed effects model revealed a significant negative association between mother’s postpartum depression score and brain signal entropy specifically within the primary somatosensory cortex (red) during affective touch (left panels) in two independent samples.

In exploratory analyses, we ran linear mixed effects models as above for each frequency band (Delta, Theta, Alpha, Beta, Gamma) and found that our results were primarily driven by entropy within the delta frequency band (Table 2, Figure 4), which is the predominant frequency band in early infancy (Louis et al., 2016).

**Figure 4.**
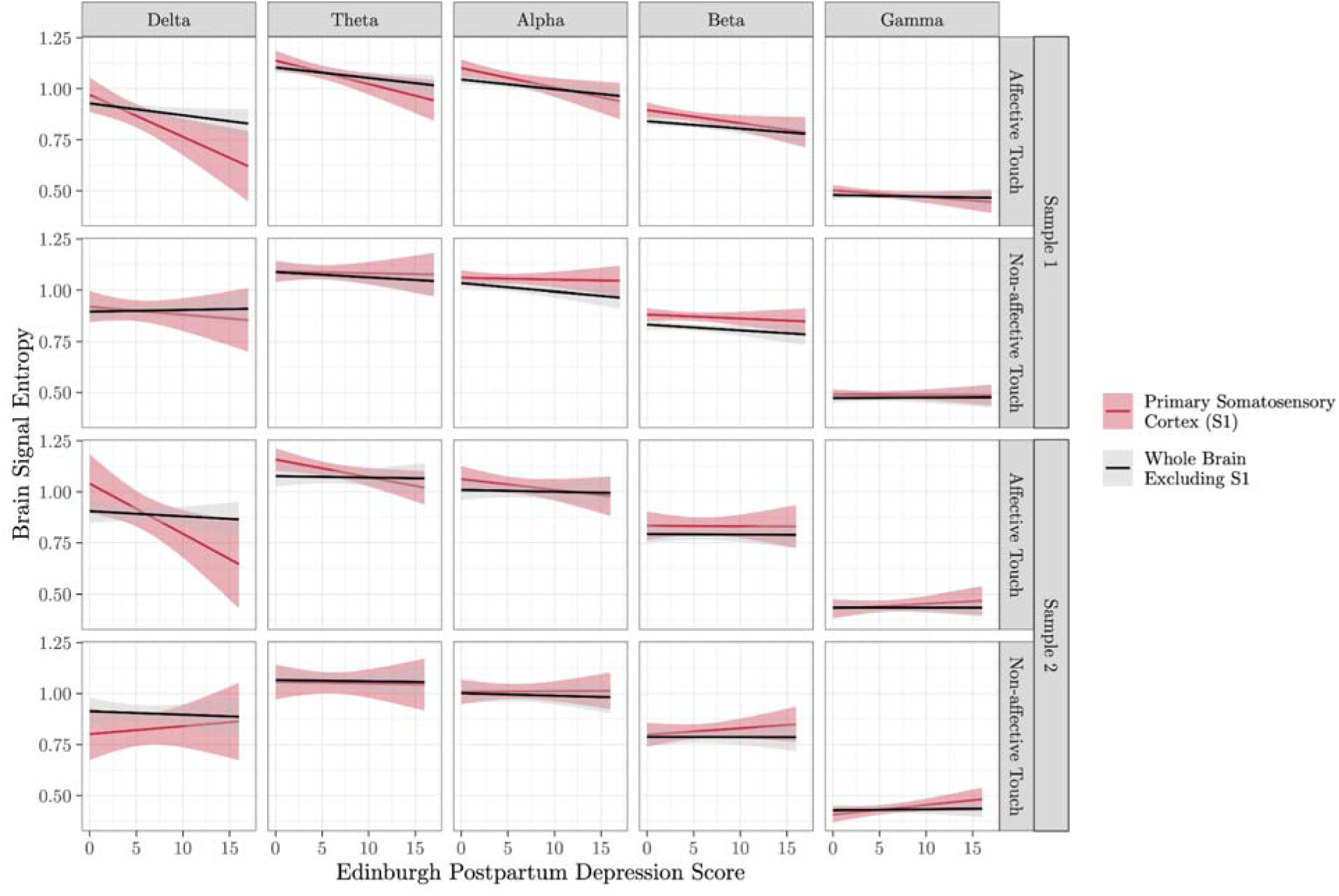
Associations between brain signal entropy and postpartum depression in primary somatosensory cortex during affective touch are driven by low frequency bands. Linear mixed effects models revealed a significant negative association between mother’s postpartum depression score and delta band (0-3 Hz) brain signal entropy within the primary somatosensory cortex (red) during affective touch (left panels) in two independent samples.

**Table 2.** Results of linear mixed effects models assessing associations between maternal postpartum depression and brain signal entropy across frequency bands. Exploratory analyses revealed significant negative associations between maternal postpartum depression and brain signal entropy in primary somatosensory cortex during affective touch were driven by entropy within the delta frequency band. S1, primary somatosensory cortex; WB, all other scalp electrodes excluding S1; *, p < .05.

| Sample | Frequency Band | Contrast | $\beta$ | p |
| --- | --- | --- | --- | --- |
| Sample 1 | Delta | S1 affective touch vs. S1 non-affective touch | 0.29 | **.012\*** |
|  |  | S1 affective touch vs. WB affective touch | 0.26 | **.026\*** |
|  |  | S1 affective touch vs. WB non-affective touch | 0.37 | **.001\*** |
|  | Theta | S1 affective touch vs. S1 non-affective Touch | 0.19 | **.008\*** |
|  |  | S1 affective touch vs. WB affective touch | 0.11 | .107 |
|  |  | S1 affective touch vs. WB non-affective touch | 0.16 | **.024\*** |
|  | Alpha | S1 affective touch vs. S1 non-affective touch | 0.15 | **.004\*** |
|  |  | S1 affective touch vs. WB affective touch | 0.08 | .109 |
|  |  | S1 affective touch vs. WB non-affective touch | 0.10 | .066 |
|  | Beta | S1 affective touch vs. S1 non-affective touch | 0.08 | .076 |
|  |  | S1 affective touch vs. WB affective touch | 0.05 | .263 |
|  |  | S1 affective touch vs. WB non-affective touch | 0.06 | .149 |
|  | Gamma | S1 affective touch vs. S1 non-affective touch | 0.05 | .079 |
|  |  | S1 affective touch vs. WB affective touch | 0.04 | .157 |
|  |  | S1 affective touch vs. WB non-affective touch | 0.06 | **.048\*** |
| Sample 2 | Delta | S1 affective touch vs. S1 non-affective touch | 0.50 | **.002\*** |
|  |  | S1 affective touch vs. WB affective touch | 0.39 | **.018\*** |
|  |  | S1 affective touch vs. WB non-affective touch | 0.40 | **.014\*** |
|  | Theta | S1 affective touch vs. S1 non-affective touch | 0.14 | .080 |
|  |  | S1 affective touch vs. WB affective touch | 0.14 | .075 |
|  |  | S1 affective touch vs. WB non-affective touch | 0.14 | .073 |
|  | Alpha | S1 affective touch vs. S1 non-affective touch | 0.10 | .090 |
|  |  | S1 affective touch vs. WB affective touch | 0.07 | .183 |
|  |  | S1 affective touch vs. WB non-affective touch | 0.07 | .216 |
|  | Beta | S1 affective touch vs. S1 non-affective touch | 0.06 | .268 |
|  |  | S1 affective touch vs. WB affective touch | 0.00 | .994 |
|  |  | S1 affective touch vs. WB non-affective touch | 0.00 | .963 |
|  | Gamma | S1 affective touch vs. S1 non-affective touch | 0.04 | .339 |
|  |  | S1 affective touch vs. WB affective touch | -0.04 | .284 |
|  |  | S1 affective touch vs. WB non-affective touch | -0.03 | .405 |

In Sample 2, we further explored the effect of touch characteristics on EPDS scores and entropy values using an engineered brush to establish that our effects were not due to acute differences in the administration of touch during the paradigm. We found that mean torque magnitude was significantly different between touch conditions (t(55)=-3.12, p=.003) such that the affective touch condition (M=7.44) had lower mean torque magnitude than the non-affective touch condition (M=10.06, Figure 5A). However, mean torque magnitude was not related to EPDS scores in either the affective touch (β=-1.45, p=.273) or non-affective touch conditions (β=-0.45, p=.566, Figure 5B). Furthermore, the slope between mean torque magnitude and brain signal entropy was not significant for affective touch within S1 (β<.001, p=.955), non-affective touch within S1 (β=-0.02, p=.446), affective touch within WB (β=-0.01, p=.609), or non-affective touch within WB (β=-0.01, p=.652, Figure 5C).

**Figure 5.**
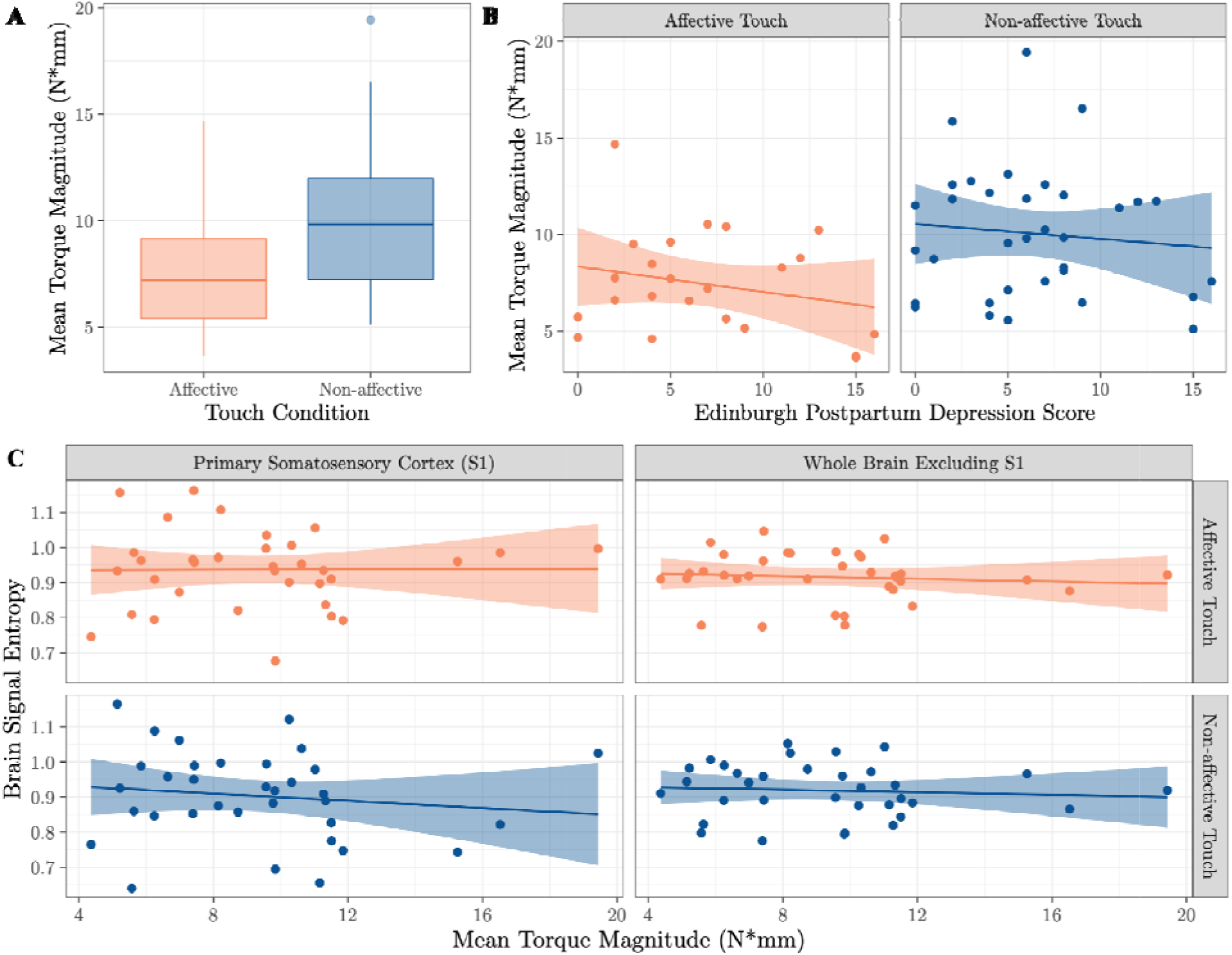
Acute touch characteristics during experimentation are unrelated to postpartum depression scores and brain signal entropy. **(A)** While mean torque magnitude was significantly lower in the affective touch (orange) relative to the non-affective touch (blue) condition, this mean torque magnitude was not related to **(B)** maternal postpartum depression scores nor **(C)** brain signal entropy in either brain region of interest.

## 5. Discussion

Here we showed in two independent samples that increased levels of maternal postpartum depression were associated with diminished infant perceptual sensitivity – i.e. lower brain signal entropy – to affective tactile stimulation specifically within the primary somatosensory cortex (S1). We further demonstrated through the use of an engineered brush that these associations were not due to acute differences in the administration of touch during the paradigm. Rather, these effects were likely reflective of chronic differences in physical contact experienced by infants related to the severity of their mother’s postpartum depression symptoms. It is well evidenced that postpartum depression influences the amount and quality of affective touch between a mother and her infant (Ferber et al., 2008; Field, 2010; Herrera et al., 2004; Malphurs et al., 1996; Mantis et al., 2019). Our results therefore suggest a link between the early caregiving environment and neural sensitivity to affective touch which is reflective of differential development of experience-dependent neural networks and an infant’s capability of forming secure attachments to their caregivers.

Our study provides a novel take on classic attachment theory by emphasizing the importance of the infant’s neural sensitivity to the early caregiving environment. Multiscale entropy quantifies the brain’s dynamic variability on multiple time scales and is reflective of perceptual sensitivity, complexity of processing, and the formation of neural networks (Duhn, 2010; Malins et al., 2018; McIntosh et al., 2008; Puglia et al., 2020; Waschke et al., 2017). Lower levels of entropy are associated with decreased perceptual sensitivity to social information and poorer social outcomes in infants (Puglia et al., 2020). We hypothesize that because babies with mothers experiencing higher levels of PPD have less chronic early life exposure to affective touch, these infants lack the experience necessary for the formation of robust neural networks that support the discrimination, processing, and interpretation of affective touch as an important social cue. This diminished experience may blunt the infant’s ability to incorporate affective touch into a complex representation of the attachment bond. We believe this poorly integrated neural network is reflected in lower levels of brain signal entropy in S1 for these infants.

S1 is a critical node in the network responsible for processing affective touch. In adults and older children, the affective properties of touch are first processed in the insula, and then go on to modulate cortical responses in S1 (Suvilehto et al., 2021). Soft brush stroking to elicit C-tactile afferents, such as that employed in the present study, has been shown to activate S1 (Liljencrantz & Olausson, 2014). Activation in this region is associated with the encoding of naturalistic social interactions. For example, S1 is capable of discriminating between touch from an attachment figure vs. a stranger (Gazzola et al., 2012; Suvilehto et al., 2021). In infants, S1 plays an even more predominant role in processing affective touch as increasing evidence suggests that the socioemotional network is not yet activated in response to affective touch (Miguel, Gonçalves, et al., 2019; Miguel, Lisboa, et al., 2019; Pirazzoli et al., 2019) until perhaps two years of age (Maria et al., 2022). Our exploratory, post-hoc investigation into the impact of age on our results is in alignment with the idea that a larger socioemotional network comes online for processing affective touch stimuli throughout development; while we replicated the same pattern of results in both younger and older infants, the difference in slope between maternal postpartum depression and brain signal entropy during affective touch within S1 vs. whole brain was less pronounced in older infants, suggesting these older infants are starting to activate a broader network of regions in response to affective touch stimuli (Figure S7).

Our results have the potential to inform clinical practice to improve the formation of strong attachment relationships for mothers suffering from postpartum depression. For instance, infants of mothers with postpartum depression often experience cognitive and motor delays (Field, 1998; Lubotzky-Gete et al., 2021); differences in neural entropy observed in this study could serve as a biomarker of these future delays allowing for early intervention and exposure to affective touch. Preterm infants and their mothers who experience a high degree of separation and high levels of maternal depression (Ozdil, 2023) may particularly benefit from education and intervention on affective touch as a mechanism for strengthening attachment and improving health outcomes (Yoshida & Funato, 2021). Touch interventions are increasingly employed in neonatal intensive care environments to improve health and developmental outcomes for preterm babies (Ferber et al., 2005). These interventions have also been shown to benefit mothers with PPD by improving mother-infant relationships and attachment (Lindensmith, 2018). Brain signal entropy may be a mechanism through which these affective touch interventions impart their health and relational benefits.

### 5.1 Limitations and Future Directions

One limitation to the present study is that we did not quantify touch between the mother-infant dyads in our samples to confirm that the common finding that mothers with postpartum depression provide less affective touch to their infant holds true in our sample. Additionally, our samples are predominately white, highly educated, and report higher income than the national average (Guzman & Kollar, 2022) (Table 1). While we replicated our results in two independent samples, future work should investigate these associations in more diverse and high-risk populations while including direct measures of mother-infant affective touch and attachment.

Our use of EEG allowed us to assess moment-to-moment variability in cortical signaling with high temporal resolution. However, given the low spatial resolution of EEG, we were unable to assess the activation of and connectivity between subcortical regions associated with affective touch processing such as the insular cortex. Future research employing MRI or dual EEG/MRI recordings will provide a more comprehensive understanding of how early life experience modulates the formation of neural networks supporting affective touch processing.

Finally, future work would also benefit from including epigenetic measures to understand how biological predisposition and variation in the early care environment interact to shape the development of experience-dependent neural networks supporting affective touch processing. For example, Murgatroyd et al (Murgatroyd et al., 2015) identified a positive association between infant glucocorticoid receptor gene (*NR3C1*) methylation and maternal postpartum depression scores, and a negative association between infant *NR3C1* methylation and maternal stroking, suggesting the infant is biologically sensitivity to maternal behavior. The oxytocinergic system is also particularly well suited to this type of investigation as it is activated during affective touch (Tsuji et al., 2015; Uvnäs-Moberg & Petersson, 2010; Vittner et al., 2018). DNA methylation of the oxytocin receptor gene (*OXTR*) has previously been linked to individual differences in brain signal entropy, influencing the salience of and sensitivity to social information (Puglia et al., 2020). Future work should investigate whether this epigenetically regulated brain signal variability impacts the experience of and neurodevelopmental response to affective touch. These investigations will inform how epigenetic modifications impact both mother’s psychopathology and infant’s sensitivity to maternal cues to drive successful mother-infant attachment.

### 5.2 Conclusion

Our combined use of techniques within developmental psychology, cognitive neuroscience, and mechanical engineering provides an innovative approach to detecting associations between postpartum depression and the formation of experience-dependent neural networks that process affective touch and support attachment formation. Attachment between mother and infant reflects an innately dyadic relationship highly dependent on both the mother’s ability to deliver affective touch appropriately as well as the infant’s perceptual sensitivity to this affective touch. Postpartum depression is associated with chronic differences in early life physical contact; these changes to the infant’s early life sensory environment shape the formation of experience-dependent neural networks. We demonstrated that the severity of maternal postpartum depression symptoms is associated with lower levels of neural entropy in infancy during affective touch. These findings suggest a mechanism by which postpartum depression influences the development of the infant’s lifelong social attachments and identifies potential neurobiological targets for intervention.

## 6. Acknowledgements

We thank the parents and infants who took part in this study and the research assistants from the Developmental Neuroanalytics Lab who assisted in collecting the data. Funding for this work was provided by the NIMH (K01MH125173) and the Jefferson Trust Foundation to M.H.P.

## 7. Author Contributions

M.G.N. and M.H.P. conceptualized the study. All authors developed the methodology. Z.T.L. and G.J.G. developed the engineered brush. M.G.N., Z.T.L., and M.H.P. collected and analyzed the data. G.J.G. and M.H.P. provided project supervision. M.G.N, Z.T.L., and M.H.P. wrote the paper. All authors edited and approved the final draft.

