## Supplemental Materials for "Infant neural sensitivity to affective touch is associated with maternal postpartum depression"

### 1. Supplementary Data

We investigated associations between Edinburgh Postpartum Depression Scale (EPDS) scores and demographic information in the combined sample for a total n = 115. Demographics for each mother were obtained through self-report questionnaire data provided during the study appointment or extracted from the University of Virginia (UVA) electronic health record. The mother's age, race, ethnicity, total household income, and highest level of education are reported in Table 1 of the main manuscript text. Of the 115 dyads, the majority were white (n = 89), and the next highest racial majority were Black or African American (n =15). Studies have consistently shown that Black and Hispanic women are more likely to develop and/or report postpartum depressive symptoms (Howell et al., 2005; Silverman et al., 2017; Onyewuenyi et al., 2023). Additional socioeconomic factors previously associated with postpartum depression include low income, having less than a college education, and being unemployed; one study found that women with these socioeconomic disadvantages were 11 times more likely to have scores indicative of postpartum depression three months postpartum (Goyal et al., 2010).

To test for associations between EPDS and demographic characteristics of the present sample, we first performed a one-way ANOVA to assess the effect of race on EPDS scores. The analysis revealed a statistically significant difference in mean EPDS score across racial groups (F(3,111) = 2.72, p = .0.48) (Figure S1). Post hoc comparisons using the Tukey Honest Significant Differences test indicated that EPDS scores only differed between Other and White racial groups (p=.031). We then performed a two-sample t-test to compare mean EPDS scores across ethnicities. We found a significant difference in ethnicity ((t(113) = 2.88, p = .005) such that EPDS scores were higher for Spanish/Hispanic women (M=8.80) compared to Non Spanish/Hispanic women (M=4.75) (Figure S2).

Next, we performed linear regressions to test for associations between EPDS scores and (1) total household income, and (2) mother’s age. We found a statistically significant negative association between household income and EPDS scores (β = -0.27, p = .005) (Figure S3). It is worth noting that the mean income in this study was $104,470.30 which is around $30,000 more than the mean income in the United States (Gloria Guzman & Melissa Kollar, 2022). Mother’s age was not significantly associated with EPDS scores (β = -0.01 , p = .570) (Figure S4). We performed a one-way ANOVA to determine the effect of education on EPDS scores. There was no statistically significant difference between groups (F(5,105)=2.01, p = .083) (Figure S5).

Next, we tested for associations between EPDS scores and birth method using a two-sample t-test. We find no significant difference in EPDS scores (t(113) = 0.49, p= .624) for women who gave birth via vaginal delivery (M= 5.24) compared to those who gave birth via caesarean delivery (M= 4.81) (Figure S6).

Finally, we conducted a post-hoc analysis to determine whether we find an age effect in the association between PPD and infant brain signal entropy. Infants less than or equal to the median age (57 days) were classified as younger infants (n=58, M age=32.64 days, SD = 12.76), and infants greater than the median age were classified as older infants (n=57, M age=105.81 days, SD = 30.62 ). We ran linear mixed effects models in R (R Core Team, 2020) using the lmerTest package (Kuznetsova et al., 2017) including touch condition (affective; non-affective), brain region of interest (primary somatosensory cortex – S1; whole brain – WB), EPDS score, and the interactions among these terms as independent variables and brain signal entropy as the dependent variable. In the younger infant sample, we found the slope between EPDS and brain signal entropy during affective touch within S1 significantly differed from the slope between EPDS and brain signal entropy during affective touch within WB (β = 0.36, p = .026), non-affective touch within S1 (β = 0.35, p = .030), and non-affective touch within WB (β = 0.44, p = .006, Figure S7). We find these associations between EPDS and brain signal entropy are maintained within the older infant sample. The slope between EPDS and brain signal entropy during affective touch within S1 differed from the slope between EPDS and brain signal entropy during affective touch within WB (β **=** 0.27, p = .066), non-affective touch within S1 (β = 0.46, p = .002), and non-affective touch within WB (β = 0.34, p = .019, Figure S7).

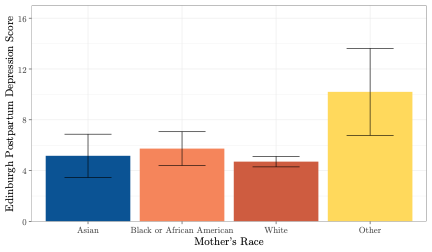

**Figure S1: Impact of race on postpartum depression scores.** A one-way ANOVA revealed a statistically significant difference in EPDS score based on race such that EPDS significantly varied across the White and Other race categories.

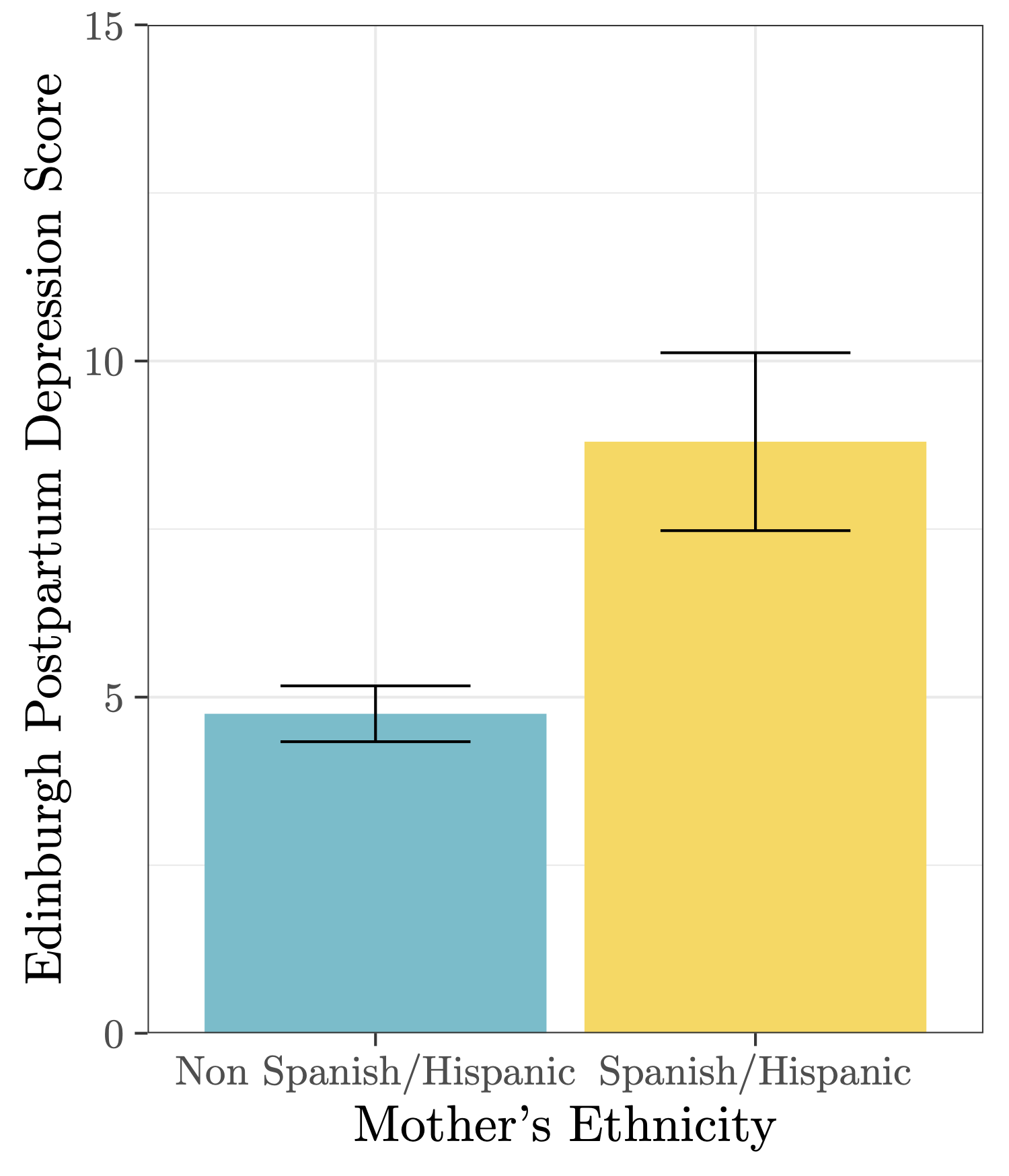

**Figure S2: Impact of ethnicity on postpartum depression scores.** A two-sample t-test revealed a significant difference in mean postpartum depression levels based on ethnicity such that Spanish/Hispanic women display higher postpartum depression levels than Non Spanish / Hispanic women.

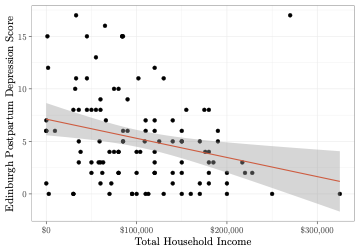

**Figure S3: Impact of total household income on postpartum depression scores.** A linear regression model revealed a significant negative association between income and postpartum depression scores.

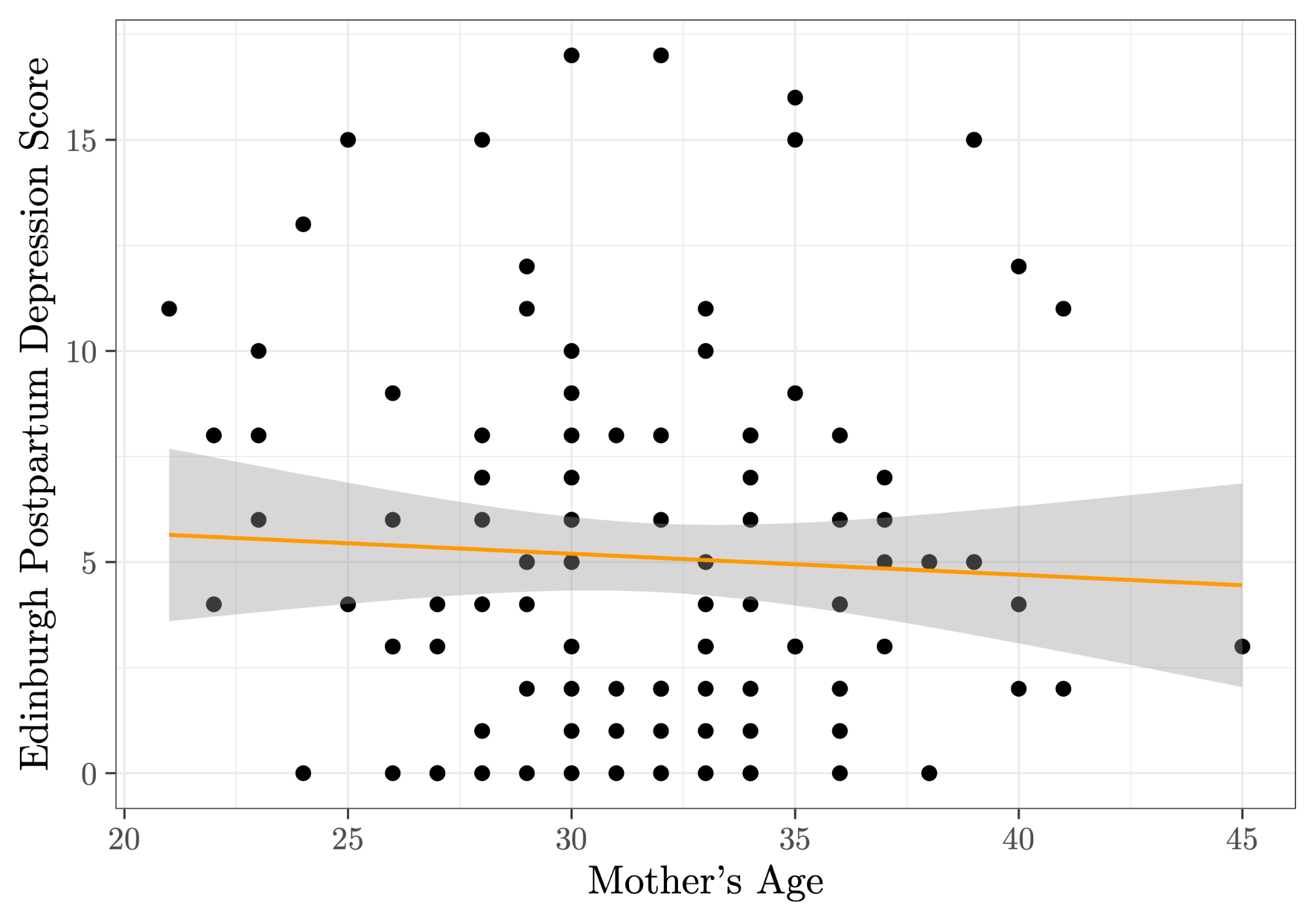

**Figure S4: Impact of mother’s age on postpartum depression scores.** A linear regression model revealed no significant association between mother’s age and postpartum depression scores.

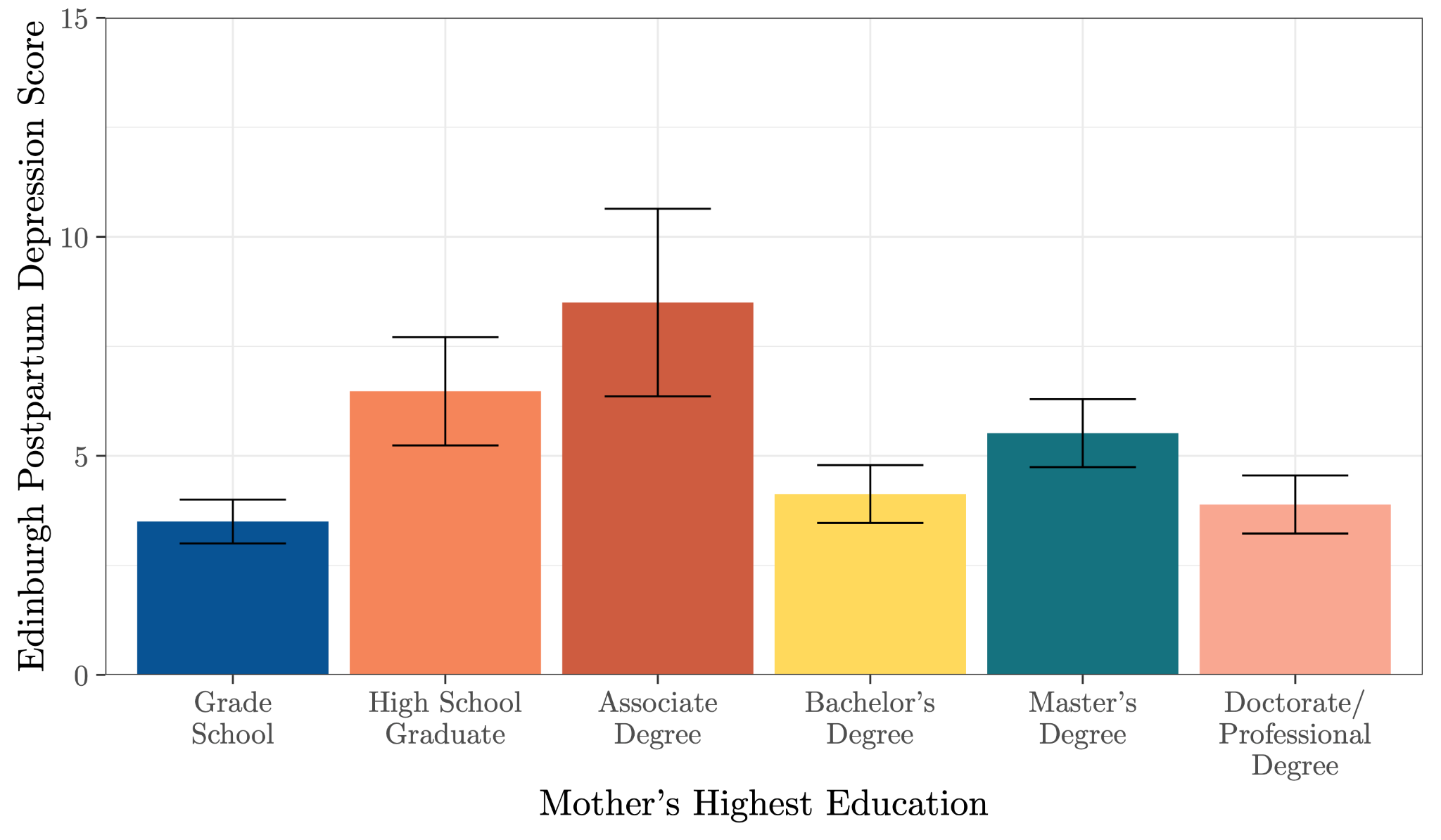

**Figure S5: Impact of highest level of education on postpartum depression scores.** A one-way ANOVA test revealed no statistically significant difference between any of the education groups.

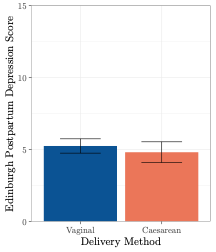

**Figure S6: Impact of delivery method on postpartum depression scores.** A two sample t-test revealed no significant difference in mean postpartum depression levels based on delivery method.

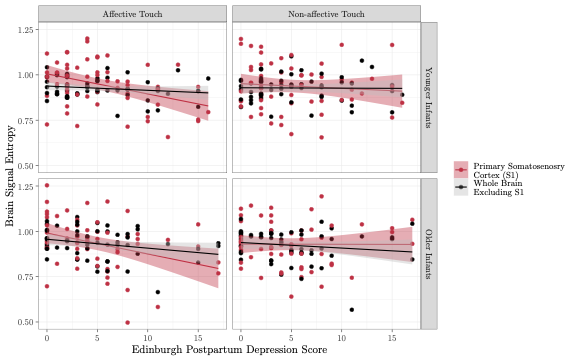

**Figure S7. Associations between brain signal entropy and postpartum depression are maintained regardless of infant age.** Infants were grouped into younger (n=58) and older (n=57) infant groups by median split. A linear mixed effects model revealed a significant negative association between mother’s postpartum depression score and brain signal entropy specifically within the primary somatosensory cortex (red) during affective touch (left panels) in both younger (top) and older (bottom) infants.

### 2. References - Supplement

Goyal, D., Gay, C., & Lee, K. A. (2010). How much does Low Socioeconomic Status Increase the Risk of Prenatal and Postpartum Depressive Symptoms in First Time Mothers? *Women’s Health Issues : Official Publication of the Jacobs Institute of Women’s Health*, *20*(2), 96–104. <https://doi.org/10.1016/j.whi.2009.11.003>

Howell, E. A., Mora, P. A., Horowitz, C. R., & Leventhal, H. (2005). Racial and Ethnic Differences in Factors Associated With Early Postpartum Depressive Symptoms. *Obstetrics and Gynecology*, *105*(6), 1442–1450. <https://doi.org/10.1097/01.AOG.0000164050.34126.37>

R Core Team. (2020). *R. Core Team, 2023. R: A language and environment for statistical computing.* [Computer software]. R Foundation for Statistical Computing.

Silverman, M. E., Reichenberg, A., Savitz, D. A., Cnattingius, S., Lichtenstein, P., Hultman, C. M., Larsson, H., & Sandin, S. (2017). The Risk Factors for Postpartum Depression: A Population Based Study. *Depression and Anxiety*, *34*(2), 178–187. https://doi.org/10.1002/da.22597

Onyewuenyi T.L., Peterman, K., Zaritsky, E, Ritterman, M.L., Weintraub, B.L. Pettway, C.P. Quesenberry, N.N., Surmava, A. & Avalos, L.A. (2023). Neighborhood Disadvantage, Race and Ethnicity, and Postpartum Depression*. Equity, Diversity, and Inclusion*, *6*(11):e2342398. doi:10.1001/jamanetworkopen.2023.42398.
